# Testing for a wildlife reservoir of divergent SARS-CoV-2 in white-tailed deer

**DOI:** 10.1101/2025.05.22.655348

**Authors:** Lauren Crawshaw, Jonathon D. Kotwa, Simon P. Jeeves, Courtney Loomer, Antonia Dibernardo, Ashley Stewart, Sasha L. Newar, Emily Chien, Winfield Yim, Peter Kruczkiewicz, Oksana Vernygora, Oliver Lung, Albrecht I. Schulte-Hostedde, Larissa Nituch, Finlay Maguire, Bradley Pickering, Claire M. Jardine, Heather Coatsworth, Samira Mubareka, Jeff Bowman

## Abstract

The 2021 discovery of a divergent lineage (B.1.641) of severe acute respiratory syndrome coronavirus 2 (SARS-CoV-2) in white-tailed deer (*Odocoileus virginianus*) from Ontario raised concerns that deer were a potential reservoir. To assess whether white-tailed deer continued to be infected with B.1.641 and to test for spillover into other species, we established a surveillance program in Ontario by sampling wildlife via existing monitoring programs and through active surveillance of captive and wild animals. Between 2022 to 2024, we tested 2,839 animals, identifying one active SARS-CoV-2 infection (a likely spillover of a recombinant XBB.2.3.11.3 lineage), but no cases of B.1.641. Overall, 93 animals (6.8%) tested positive for SARS-CoV-2 antibodies, including 89 white-tailed deer, two Virginia opossums (*Didelphis virginiana*), one American mink (*Neogale vison)*, and one river otter (*Lontra canadensis*). In Southwestern Ontario, where B.1.641 was originally detected, 15.2% of deer samples were seropositive. Generalized Linear Models demonstrated that seropositive deer were more likely to be found in areas with a higher fall deer harvest and human population density, and closer to previous B.1.641 cases. Our data suggest that deer-associated B.1.641 may have caused a relatively localized epizootic without forming a stable reservoir. This study underscores the importance of One Health-focused surveillance.

## Introduction

Severe acute respiratory syndrome coronavirus 2 (SARS-CoV-2) is responsible for coronavirus disease 2019 (COVID-19) and an estimated 7 million human deaths worldwide (World Health Organization 2024). SARS-CoV-2 continues to be a major public health concern post-pandemic, and the effectiveness of human-centred interventions, such as vaccination, risk being undermined by a variant persisting outside of human populations. When a pathogen is able to infect multiple host species, it is more likely to be an emerging human pathogen (Singh et al. 2023). Understanding which species are susceptible to SARS-CoV-2 can shed light on how the virus is able to spill into humans (Palmer et al. 2021). SARS-CoV-2 has a broad host range, infecting species from almost all orders of mammal (Abdel-Moneim and Abdelwhab 2020; Zhou et al. 2020; Kuchipudi et al. 2023; Z.H. Tan et al. 2024). The propensity of SARS-CoV-2 to infect numerous species can be partially explained by the virus’ ability to bind to the angiotensin-converting enzyme 2 (ACE2) host cell receptor (F. Wu et al. 2020), which is conserved across many mammalian species (Damas et al. 2020; Zhou et al. 2020).

Given the potential for SARS-CoV-2 spillback from humans to animal populations, there remains a pressing need to understand the risk of the virus infecting novel species (C.C.S. Tan et al. 2024). Transmission of SARS-CoV-2 from humans into other mammalian species occurred early in the pandemic. The first human-to-animal transmission events of SARS-CoV-2 occurred in mink farms of the Netherlands and Denmark in 2020, likely transmitted from farm workers (Oreshkova et al. 2020; Oude Munnink et al. 2021; Wolters et al. 2022). Since then, there have been many reports of infected animal populations, from both domesticated and wild species. For example, in a study of wildlife in Virginia, USA, 8 of 23 native mammalian species tested positive or seropositive. SARS-CoV-2 infection was strongly correlated with urbanization and prevalence was high in synanthropic species such as Virginia opossums (*Didelphis virginiana*) and raccoons (*Procyon lotor*) (Goldberg et al. 2024). Infection of 11 cattle in Germany coincided with a large wave of the Delta variant of concern in humans in fall to early winter 2021 (Wernike et al. 2022). In Nigeria, SARS-CoV-2 antibodies were detected in feral dogs, rabbits, and pangolins (Agusi et al. 2024), in the context of an estimated 80% seroprevalence among humans (Kolawole et al. 2022).

The broad host range of SARS-CoV-2 and frequent spillback from humans to wildlife has led to concern regarding the establishment of a potential reservoir. We follow the suggestion of Haydon et al. (2002) and define a reservoir as having two characteristics: an ability to maintain infection with a zoonotic pathogen, and the presence of an interface with humans. Reservoirs may undermine public health efforts and make strain-specific vaccination more challenging by providing a continual source of virus capable of transmitting back into human populations (Haydon et al. 2002), gaining genomic diversity through recombination with other variants (Wei et al. 2021) or adaptive immune-evasive mutations. Reservoirs have the potential to spread a pathogen back to vulnerable human populations even after the pathogen has been reduced or eradicated (Haydon et al. 2002). Such is the case with Middle East respiratory syndrome (MERS-CoV), a betacoronavirus that first emerged in 2012 in Jordan and Saudi Arabia. While MERS outbreaks in human populations are typically self limiting (Pustake et al. 2022), the pathogen is widely enzootic in dromedary camels (Mackay and Arden 2015), with camel-to-camel transmission exclusively perpetuating the epidemic and evolution of the virus (Dudas et al. 2018).

With the concern about SARS-CoV-2 spillback into wildlife, many jurisdictions in North America initiated surveillance early in the pandemic. White-tailed deer (*Odocoileus virginianus,* hereafter deer) were flagged as an important species for surveillance, given their abundance, highly social and transient behaviour (Peterson et al. 2017), frequent contact with humans (Pratt et al. 2025), and high affinity for ACE2 virus binding (Damas et al. 2020). Challenge studies found that deer fawns are not only highly susceptible to SARS-CoV-2, but they could transmit the infection to other fawns through droplets or aerosol (Palmer et al. 2021; Martins et al. 2022), and the virus can be transmitted vertically from doe to fetus (Cool et al. 2022). Surveillance studies found infected deer in wild free-ranging herds in 23 US states (Feng et al. 2023), with dozens of spillover events from humans and evidence of sustained and continual deer-to-deer transmission (Hale et al. 2022; Kuchipudi et al. 2022; Marques et al. 2022; Pickering et al. 2022; Caserta et al. 2023; Feng et al. 2023; McBride et al. 2023). SARS-CoV-2 antibodies were detected in almost half the deer sampled (40%) across four US states in 2021, with the highest prevalence in Michigan (67%) (Chandler et al. 2021). Reports have shown the circulation of nearly extinct variants of concern like Alpha or Gamma (or both) persisting in deer many months after their last detection in humans in the US population (Marques et al. 2022; Caserta et al. 2023; Feng et al. 2023; McBride et al. 2023).

Recognizing the potential for transmission of SARS-CoV-2 from humans to deer, we initiated surveillance of free-ranging deer harvested by hunters in southern Ontario, Canada. Surveys in autumn 2021 produced evidence of sustained deer-to-deer transmission within 80 km of Michigan, and a divergent lineage of SARS-CoV-2 infecting deer, Phylogenetic Assignment of Named Global Outbreak (PANGO) lineage B.1.641. This lineage contains 76 nucleotide mutations relative to ancestral SARS-CoV-2 (Wuhan Hu.1) and 49 mutations compared to its closest ancestor in Global Initiative on Sharing All Influenza Data (GISAID) as of March 2022, making this one of the most divergent SARS-CoV-2 lineages sequenced at the time of discovery. B.1.641 shares a common ancestor with human- and mink-derived sequences from Michigan sampled one year prior (autumn 2020). The B.1.641 lineage also stands out for being the first confirmed case of deer-to-human transmission documented up to that point, making it the first confirmed wildlife-to-human spillover event of SARS-CoV-2. The associated human case (ON-PHL-21-44225), which shared the majority (80/90) of mutations with the deer samples, was detected in autumn 2021 in a person with known contact with deer a week prior to symptom onset (Pickering et al. 2022).

Viruses that evolve in the same species show similar mutational spectra due to selective pressure from the host environment (Shan et al. 2020; Wei et al. 2021; Rudar et al. 2024). Analysis of mutational spectra in B.1.641, especially the prevalence of C-to-U base pair changes, suggest that this virus evolved for some time in deer between 2020 and 2021. Consequently, we infer that this lineage is deer-adapted, and has rapidly accumulated mutations, suggesting the virus was able to propagate freely in a highly susceptible host population (Pickering et al. 2022). Given the possibility of sustained infection of deer with B.1.641, we were interested in the potential establishment of a deer reservoir of SARS-CoV-2, and the role that a deer host might play in altering the evolutionary trajectory of this lineage. Mutations in the spike protein account for nine of the 76 mutations of B.1.641 (Pickering et al. 2022). Spike protein mutations can improve affinity for ACE2 and expand the host range (Oreshkova et al. 2020; Wei et al. 2021; McBride et al. 2023; Wang et al. 2023; Goldberg et al. 2024). Among US states with infected deer, the virus has shown repeated adaptation to the deer host through substitutions in the spike protein and other proteins (Feng et al. 2023). The potential for deer-adapted mutations to increase risk to humans remains a concern.

In response to the potential establishment of a reservoir of B.1.641 in deer in Ontario, we initiated a surveillance program to sample a wide array of captive and wild animal species across the same general area where the B.1.641 lineage had been discovered. We carried out active surveillance in southern Ontario across a suite of mammal species, with a specific focus on deer given the potential for reservoir establishment in this species. We examined the role of deer as a host and reservoir, including whether B.1.641 has been sustained in deer or transmitted into other species. In anticipation of finding more infected deer, we planned to sequence viral genomes, aiming to characterize virus mutational spectra and build our understanding of the host response. We hypothesized that sustained infection of deer with B.1.641 would lead to viral genomes in deer closely related to B.1.641, rather than infection with lineages more commonly circulating in the human population at the time of the study. Likewise, spillover of B.1.641 from deer to other animal species would lead to the discovery of lineages closely related to B.1.641.

## Materials & methods

### Sample collection

Between November 2019 and June 2024, we collected mid-turbinate, nasopharyngeal, or oral swabs and blood samples from live and deceased animals across southern Ontario. Samples were sourced through live-trapping, roadkill collection, existing ecological monitoring projects, and from captive facilities (i.e. wildlife rehabilitation facilities, private zoos). Sampling priority was given to species known to be susceptible to SARS-CoV-2, including cervids (Hale et al. 2022; Kuchipudi et al. 2022), mink (*Neogale vison)* (Oude Munnink et al. 2021), deer mice (*Peromyscus maniculatus*), and white-footed mice (*Peromyscus leucopus*) (Bosco-Lauth et al. 2021; Fagre et al. 2021; Griffin et al. 2021) as well as species suspected to be susceptible (e.g. canids, Virginia opossums, mustelids). We also opportunistically sampled mammalian species as they were available, including those with unknown or limited evidence of susceptibility (e.g. beaver, *Castor canadensis*). Two sizes of sterile swabs (Hardy Diagnostics™ Flex Minitip Nasopharyngeal Flock Swab, Puritan™ PurFlock™ Ultra Flocked Swabs) (Thermo Fisher Scientific Inc; https://www.fishersci.ca) were used to collect oral or nasal swabs, with the minitip swabs used to collect oral swabs from animals the size of *Peromyscus*.

#### Wildlife rehabilitators and zoos

Mid-turbinate or oral swabs were collected from captive animals at 11 registered wildlife rehabilitators and five regional zoos. The type of swab collected depended on the species, size, and ability to humanely access the nasal passage. To limit the demand on the facility and number of samples, only one or two animals were tested from litters of juveniles, and neonates were excluded. While most of the samples were collected by the visiting research personnel, six facilities were also provided with sampling kits and a protocol to collect additional samples.

#### Pre-existing surveillance and ecology programs

Most samples were collected through pre-existing wildlife projects, with the largest source being deceased deer tested through Ontario’s Chronic Wasting Disease (CWD) surveillance program, as described by Pickering et al. (2022). CWD testing is carried out each November and December in a subset of administrative areas within Ontario, known as Wildlife Management Units (WMUs). Each year a handful of WMUs are selected for CWD surveillance based on a provincial CWD risk model (Ontario Ministry of Natural Resources 2019). Every deer is assigned to one 10 x 10 km square location where it was harvested, which corresponds to a vector grid layer sourced from the Ontario Breeding Bird Atlas (Birds Canada 2024). Since 2021, SARS-CoV-2 molecular detection and serology have been incorporated into the province’s seasonal CWD project through the collection of nasopharyngeal swabs and blood samples. Archival retropharyngeal lymph nodes (RPLN) and blood collected in 2019 and 2020 were also retrospectively tested.

Opportunistic sampling for SARS-CoV-2 took place during ongoing ecological studies run by the Ontario Ministry of Natural Resources (MNR) and Trent University. During the MNR’s rabies vaccine serosurveillance program, SARS-CoV-2 swabs and blood were collected from target species (i.e., raccoons and striped skunks, *Mephitis mephitis*) in the fall of 2020 and 2021, and from any mink or fox bycatch during their trap-vaccinate-release (TVR) project in the summer and fall of 2022, 2023, and 2024. Between 2022 and 2024, other projects involved in collecting SARS-CoV-2 samples included projects placing GPS transmitters on eastern wild turkeys *(Meleagris gallopavo silvestris*), deer, moose (*Alces alces)*, the Algonquin wolf (*Canis lupus lycaon),* and eastern coyote (*Canis latrans*), as well as wild boar removed from the landscape (*Sus scrofa*). The majority of *Peromyscus* samples came from trap lines in Algonquin Provincial Park as part of the Algonquin Small Mammal Project (Fryxell et al. 1998).

#### Roadkill and carcasses

From 2022 to early 2024, swabs and blood were collected from any fresh target species found dead on roadsides during live-trapping work (i.e., deer, skunk, red fox, mink). This also included any deer roadkill picked up and reported by participating municipal patrol lots. When possible, blood was collected directly from the heart muscle. Four deceased deer were sampled for SARS-CoV-2 after strange behaviour prior to death, two of which tested positive for epizootic hemorrhagic disease. Additionally, local trappers provided carcasses of mink and other furbearers (beaver (*C. canadensis*), fisher (*Pekania pennanti*), river otter (*Lontra canadensis*)) between 2021 and 2023 where tissue samples, swabs, and blood were collected for SARS-CoV-2 testing.

#### Live-trapping study area and trapping risk model

We carried out active surveillance through live-trapping of small mammals and mesocarnivores. Given the geographic expanse of southern Ontario, the challenge in accessing wildlife hosts, and the low prevalence of SARS-CoV-2 up to May 2021 (Greenhorn et al. 2022), we built an ecological model to predict where humans, wildlife, and the virus were more likely to be in contact (Kuchipudi et al. 2023), which narrowed the geographic scope for active surveillance. Our model included: (1) proximity to the 17 SARS-CoV-2 B.1.641 PCR-positive deer from 2021 (Pickering et al. 2022); (2) approximate distance to mink farms and (3) deer farms; (4) distance to the state of Michigan (U.S. Census Bureau 2018); and (5) an independently derived estimate of terrestrial landscape connectivity referred to as “current density” (Pither et al. 2023).

Spatial analyses were carried out using ArcGIS Pro 3.0.3 (Esri 2022). Euclidean distance was calculated for the three vector-based point layers (i.e., deer and mink farms, deer PCR cases) to the same resolution and coordinate system. Pixel values of every raster (100 m resolution) were then reclassified to a scale from 10 to 1, with higher scores representing higher potential risk of SARS-CoV-2 transmission (e.g., the closer to a deer or mink farm, the higher the score). The spatial grid vector layer (Birds Canada 2024) used for CWD surveillance was used as the sampling unit, and mean zonal statistics were calculated for each grid cell for all five variables. Each grid cell was then averaged to obtain a mean risk score, with proximity to the 17 PCR-positive deer given double the weight as the other variables, given our priority to sample near previous B.1.641 positive cases:

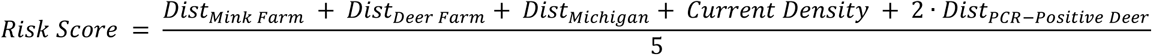

The results from the risk model identified an area with the highest risk score (mean score >8), covering 17,000 km^2^, which included the metropolitan area of the City of London, Ontario. This risk area was prioritized for sample collection via live-trapping. The trapping zone included a Komoka Provincial Park and six provincial green space management areas, called Conservation Authorities, which intersected the Thames River waterways.

#### Live-trapping surveys

Live trapping was carried out across a 5,194 km² region of southern Ontario over three years (June - October 2022 and 2023, one week in February 2024). Many trapping sites from 2022 were revisited in 2023. During the trapping period, small mammals (i.e., rodents, shrews) and mesocarnivores (i.e., raccoons, skunks, opossum, mink) were live-trapped on private land and government managed greenspace, with permission obtained through informed consent by landowners, parks, and conservation authorities. Various sizes of Tomahawk live trap (sizes 102, 103, 105, and 106) (Tomahawk Live Trap; https://www.livetrap.com) were baited with sardines and scent lures. Small mammals and shrews were trapped using Sherman live traps (25.4 cm x 7.6 cm x 7.6 cm) (H.B. Sherman Traps Inc; https://shermantraps.com) baited with a mix of sunflower seeds, peanuts, dried insects, and seafood. Each day, 40-60 Tomahawk traps and 30-40 Sherman traps were set in riparian or forest habitat for four consecutive nights per week. Traps were placed in sheltered areas for protection from the elements and secured with cables to prevent rolling. Traps were checked each morning and reset for the next day, with Sherman traps closed during the day and reset in the evenings. If evening temperatures fell below -20°C, traps would remain closed overnight. Adequate thermal conditions for trapping in the fall and winter were maintained by making sure all traps were sheltered from precipitation and wind by covering traps with plastic and supplying each trap with either cotton bedding or straw. For animals sampled on the Algonquin Small Mammal Project, Sherman and Longworth (Longworth Scientific Instrument Co; https://www.penlon.com) live traps were baited with water-soaked sunflower seeds set on established transects between May and August of each sampling year.

#### Ethics approval

Animal care and handling was approved for each year of the study under the MNR Wildlife Animal Care Committee ((WACC Protocols #23-358 and #24-491), the Trent University Animal Care Committee (#28123), and the Laurentian University Animal Care Committee (#6011106), Ontario, Canada. Live-trapping and sampling of animals was conducted in accordance with applicable laws and regulations.

#### Animal handling

To obtain samples, animals were carefully restrained in the trap or transferred into a semi-soft funnel or large plastic bag, allowing for gentle immobilization of the animal during oral swabbing and marking. Trap locations were changed every week, while the Algonquin Small Mammal Project used the same established trap lines each year. Every capture was marked with livestock paint or chalk, unless a trapping site was used for more than one consecutive week, in which case an ear tag was applied instead (sizes 1005-1 for *Peromyscus*, skunk and mink; and 1005-3 for raccoons) (National Band & Tag Company; https://www.nationalband.com). Trappers wore medical grade face masks and nitrile gloves during processing to prevent the possible spread of SARS-CoV-2, and tools (pliers, clippers) were disinfected between animals.

#### Sample collection

Swabs were gently inserted into the mouth or nasal mid-turbinates and rotated for three to five seconds. Ungulates were the only animals where it was possible to humanely swab the nasal cavity, and nasopharyngeal samples were only collected from deceased deer. A small subset of deer, mink and *Peromyscus* had an additional exterior rectal swab taken, given potential of detecting the virus in *Peromyscus* (Griffin et al. 2021), deer (Martins et al. 2022) and ferrets (*Mustela furo*) (Gortázar et al. 2021) through this route. Where possible, live-trapped mesocarnivores had a small blood sample (0.05-0.2 ml) collected on a piece of Advantec Nobuto filter paper from the tail artery or a nail clipping (Advantec MFS, Inc; https://www.advantecmfs.com). Additional tissues collected over the project included a small section of the right and left RPLNs from CWD-tested deer (2019-2020), cloacal swabs from wild turkeys, fecal samples from live-trapped deer, and lung and intestinal tissues from harvested mink.

Swab tips were stored in sterile 2 ml tubes containing 1 ml of universal transport medium (UTM; Sunnybrook Research Institute). Organs, RPLNs, and fecal material were stored in sterile dry tubes. All samples taken in the field were kept on ice packs until they could be transferred to storage at -80 °C. Blood samples collected on filter paper were allowed to air dry then placed in coin envelopes before eventually being frozen at -80 °C.

### Diagnostic testing

#### RNA extraction and PCR testing

The majority of RNA extractions were undertaken by staff at the Sunnybrook Research Institute (Toronto, Ontario, Canada) and performed using 140 μL of swab sample spiked with Armored RNA enterovirus (Asuragen; https://www.asuragen.com) using the Nuclisens EasyMag using Generic Protocol 2.0.1 (bioMérieux Canada Inc., St-Laurent, QC, Canada) according to manufacturer’s instructions. Tissue samples were thawed, weighed, minced with a scalpel, and homogenized in 600 μL of lysis buffer using the Next Advance Bullet Blender (Next Advance, Troy, NY, USA) and a 5 mm stainless steel bead at 5 m/s for 3 minutes. RNA from 30 mg tissue samples was extracted via the Nuclisens EasyMag using Specific Protocol B 2.0.1; RNA was eluted in 50 μL. RNA extractions for *Peromyscus* swab samples in UTM were performed at the Department of Pathobiology, University of Guelph (Guelph, Ontario, Canada). RNA was extracted from 140 µl of swab UTM using the QIAamp Viral RNA Mini Kit (QIAGEN; https://ww.qiagen.com) according to the manufacturer’s protocol. Extractions were spiked with 1 µl Armored RNA Enterovirus (Asuragen; https://www.asuragen.com). Final elution volume of extracted RNA was 60 µl in AVE buffer. All RT-PCR was carried out by Sunnybrook Hospital Research Institute (SRI).

A subset of certain species (i.e. mink, fox, skunk), or animals exhibiting abnormal behaviour were screened for highly pathogenic avian influenza virus (HPAIV) by the Animal Health Lab (University of Guelph, Ontario). Only HPAIV-negative samples were submitted to SRI. RT-PCR was used for SARS-CoV-2 RNA detection as previously described by Pickering et al. (2022); two gene targets were used for SARS-CoV-2 RNA detection: the 5’ untranslated region (UTR) and the envelope (E) gene (Corman et al. 2020; LeBlanc et al. 2020). All samples were run in duplicate and samples with cycle thresholds (Ct) < 40 for both SARS-CoV-2 targets and armored RNA enterovirus in at least one replicate were considered positive. A subset of *Peromyscus* spp. (N=282) samples were tested for SARS-CoV-2 RNA with a nested pancoronavirus RT-PCR assay as previously described (Kotwa et al. 2025).

#### SARS-CoV-2 antibody testing

Blood samples were shipped to the National Microbiology Laboratory in Winnipeg, Manitoba for SARS-CoV-2 antibody testing. Once received, the strips were sorted and evaluated for degree of saturation. The saturated portion of each strip was cut into 4-5 pieces using sterile scissors and dropped into a 2 ml sample tube and eluted in 1X phosphate-buffered saline (PBS) at 4°C overnight. Fully saturated (100%) strips were eluted in 0.360 ml of PBS, 75% saturated strips in 0.270 ml of PBS and 50% saturated strips in 0.180 ml of PBS. These elution volumes were considered to be equivalent to a single 10-fold dilution of serum. The eluates were tested with the Genscript SARS-CoV-2 Surrogate Virus Neutralization Test (Catalog L00847-A) (Genscript; https://www.genscript.com) following the manufacturer’s instructions. A percent inhibition greater than or equal to 30% was considered positive for the presence of SARS-CoV-2 neutralizing antibodies as per the kit protocol.

#### Whole-genome sequencing

Whole-genome sequencing on RT-PCR positive samples were conducted at SRI and was performed as follows. LunaScript RT Supermix 5X kit (New England Biolabs, NEB; https://www.neb.ca) was used to convert RNA samples to cDNA for amplification with the SARS-CoV-2 ARTIC protocol. ARTIC v4.1 10µM primer pools were used in conjunction with Q5 High-Fidelity 2X Master Mix (NEB) for the PCR reaction. Cycling conditions were as follows: 98°C for 30s, 35 cycles of 98°C for 15s, and 63°C for 5min. Post amplicon generation, the PCR products of the primer pools were combined followed by a 1X bead clean-up using AMPure XP beads (Beckman Coulter; https://www.beckmancoulter.com). Amplicon concentrations were determined using the Qubit dsDNA quantitation, High Sensitivity Kit (Thermo Fisher; https://www.thermofisher.com) on a Qubit 4 Fluorometer. Sequencing libraries were generated using the Illumina DNA prep kit as per manufacturer’s instructions. Libraries were loaded on a MiniSeq (Illumina; https://www.illumina.com) using the MiniSeq Mid Output Kit to perform paired-end sequencing (2×149bp, 300 cycles). Original material from positive samples was sent to the Canadian Food Inspection Agency (CFIA) for confirmatory RT-PCR testing.

#### Genomic analysis

Reference-mapped assembly and genomic analysis of paired-end Illumina reads was conducted with the nf-core/viralrecon Nextflow workflow (v2.6.0) (Di Tommaso et al. 2017; Patel et al. 2023). Briefly, nf-core/viralrecon ran the following steps: sequencing read quality control with FASTQC (v0.11.9) (Andrews 2010); read quality filtering and adapter trimming with fastp (v0.23.2) (Chen et al. 2018); read alignment to Wuhan-Hu-1 SARS-CoV-2 reference (MN908947.3) (L. Wu et al. 2020) with Bowtie2 (v2.4.4) (Langmead and Salzberg 2012); NEB VarSkip primer trimming from read alignments with iVar (v1.4) (Grubaugh et al. 2019); read alignment coverage and statistics calculation with Mosdepth (v0.3.3) (Pedersen and Quinlan 2018) and Samtools (v1.16.1) (Li et al. 2009); variant calling and consensus sequence generation with Bcftools (v1.16) (Danecek et al. 2021); variant effect prediction and annotation with SnpEff (v5.0) (Cingolani, Platts, et al. 2012) and SnpSift (v4.3.1t) (Cingolani, Patel, et al. 2012); PANGO lineage (Rambaut et al. 2020) assignment with Pangolin (v4.3) (O’Toole et al. 2021) using UShER (v0.6.2) (Turakhia et al. 2021) Scorpio (v0.3.17) and Constellations (v.0.1.2) (Colquhoun et al. 2023). The results from nf-core/viralrecon were summarized into an XLSX report with xlavir (v1.0.2) (https://github.com/CFIA-NCFAD/xlavir). Nextclade (v3.12.0) analysis was performed on the nf-core/viralrecon assembled consensus sequence of the PCR-positive deer (wcov1242165) using the SARS-CoV-2 “nextstrain/sars-cov-2/wuhan-hu-1/orfs” Nextclade dataset (last updated 2025-04-01 08:20:12 (UTC)) (Aksamentov et al. 2021).

#### Phylogenetic Analysis

Phylogenetic placement of the SARS-CoV-2 isolated from one positive deer was inferred relative to other publicly available sequences from white-tailed and mule deer (N=811; GISAID accessed 2025-03-19) as well as the 96 closest sequence matches recovered by UShER (dataset of 16,884,360 genomes from GISAID, GenBank, COG-UK, and CNCB (2025-03-26)).

The Wuhan-Hu-1 (GenBank accession MN908947.3) sequence was included as an outgroup. The multiple sequence alignment was performed using Nextclade v.3.12.0 and maximum likelihood phylogenetic inference analysis was performed with IQ-TREE v.2.2.3 (Minh et al., 2020). The best-fit nucleotide substitution model (GTR+F+R6) was determined by IQ-TREE ModelFinder (Kalyaanamoorthy et al. 2017) under the Bayesian Information Criterion (BIC). Node support was estimated using the ultrafast bootstrap and SH-aLRT with 1000 replicates for each test (Hoang et al. 2017). Tree visualization was performed using FigTree (http://tree.bio.ed.ac.uk/software/figtree) and Inkscape v.1.0.2 (https://inkscape.org).

### Regression modeling

Understanding the spatial patterns of infection on the landscape can shed light on the current state of SARS-CoV-2 in southern Ontario wildlife and identify risk factors for transmission. We ran regression analysis on all serological data collected since 2022 (after the discovery of the B.1.641 lineage) from deer south of latitude 47.3°N. In addition, we also explored biological characteristics of the animals to see if there was any association with past infection. All analyses were carried out in R (version 4.3.2) and RStudio (2024.04.02, Build 764) (R Core Team 2024), and mapping and graphs in R and ArcGIS (Esri 2022).

#### Zonal statistics

Landscape variables, some of which were also used in the risk model, included: (1) proximity to the 17 B.1.641 PCR-positive deer from 2021; (2) Euclidean distance to the state of Michigan (km); (3) proportion of each grid cell classified as deer/non-deer habitat (Kennedy-Slaney et al, unpublished report, 2018) derived by reclassifying the Ontario Land Cover Compilation Data (Ontario Ministry of Natural Resources 2014); (4) annual fall deer harvest in Ontario by WMU (Ontario Ministry of Natural Resources 2024); (5) current density (Pither et al. 2023); (6) average cases of human SARS-CoV-2 in the fall of each year (2020-2023) (Ontario Agency for Health Protection and Promotion (Public Health Ontario) 2024); and (7) human population (Statistics Canada 2023a) (Table 1). Due to the limited data on deer abundance in Ontario, annual deer hunting harvest was used as a proxy for deer abundance. The human population variable was created by joining the 2021 census data (Statistics Canada 2023a) to a spatial layer of the smallest census subdivisions, known as dissemination areas (Statistics Canada 2023b). Prior to running zonal statistics, the three polygon layers (human SARS-CoV-2 cases, deer harvest, and deer habitat) were converted to raster grids at a 100 m resolution. Each deer sample fell within one of the 10 x 10 km grid cells used in the risk model (Birds Canada 2024), which were used as the main sampling units to extract a zonal statistic (a weighted mean) for each of the landscape variables. In cases where the grid cells overlapped with large lakes (>5 km²), the grid cell layer was first clipped using a high resolution waterbody layer (Ontario Ministry of Natural Resources 2020).

**Table 1.** Potential variables for use in the regression models. Some variables were removed due to multicollinearity.

| Class | Variable | Description/Biological relevance |
| --- | --- | --- |
| Binary outcome | SARS-CoV-2 exposure* | Serology result (PVE+/NVE-) of each deer since January 2022. The dependent variable of interest. |
| Raster | Distance to 17 PCR+ deer* | Euclidean distance to 17 PCR+ deer of B.1.641 lineage sampled in 2021 (Pickering et al. 2022) |
| Vector to Raster | Annual fall deer harvest* | Average count of deer harvested each fall by WMU (2022-2023) (Ontario Ministry of Natural Resources 2024). Models were tested with each year, and an averaged variable containing all years. |
| Vector to Raster | Human population* | Mean human population derived from 2021 Census data (Statistics Canada 2023a) per dissemination area (Statistics Canada 2023b). |
| Vector to Raster | Human Density | Human population per square kilometre, derived from 2021 Census data (Statistics Canada 2023a) per dissemination area (Statistics Canada 2023b). |
| Vector to Raster | Human COVID-19 cases* | Daily change in human cases of COVID-19 reported to public health units each fall (September - December 2022-2023) (Ontario Agency for Health Protection and Promotion (Public Health Ontario) 2024). Models were tested with each year, and a variable containing all years. |
| Factor | Year* | Year sample was collected (2022-2024) |
| Raster | Distance to Michigan | Euclidean distance to the state of Michigan (U.S. Census Bureau 2018) |
| Raster | Deer habitat | Binary reclassification of the Ontario Land Cover Compilation Data (Ontario Ministry of Natural Resources 2014) as deer habitat or non-habitat (Kennedy-Slaney et al. 2018, unpublished) |
| Raster | Current density* | Ecological connectivity layer derived from costs from anthropogenic and natural barriers, with higher values equating lower cost for terrestrial animals to travel across the landscape (Pither et al. 2023) |

\*Variables used in final full model

#### Sampling unit selection

Prior to running the models, a second sampling unit (30 x 30 km) was created to explore possible bias of having a single 10×10 grid cell represent the presence of a highly mobile deer species with a wide potential home range, with some yearling males reported to range over 6000 hectares (Lesage et al. 2000). This was particularly important where the true deer locations occur near the edges of the 10 x 10 km grid cell. This second sampling unit was created by buffering each grid cell by 10 km. Zonal statistics were repeated with these two buffer sampling units for variables unrelated to distance and compared to the grid-based sampling unit for correlation and relationship with the dependent variable.

Zonal mean of the larger sampling unit were highly correlated with the same variable run with the grid (>0.75), indicating high similarities with the buffer and grid cell zonal statistics. Due to this similarity, plus less multicollinearity between the original 10 x 10 grid variables of interest, the original 10 x 10 km grid sampling unit was selected moving forward with the models.

#### Modeling

We used Generalized Linear Mixed Models (GLMMs) through R package *lme4* version 1.1-37 (Bates et al. 2015) to identify landscape features associated with a serological history of infection in wild deer. The 10 x 10 km grid cell was used as the random effect to account for non-independence between samples within these larger groups. Candidate models were selected by first identifying highly correlated variables and retaining the variable that were perceived as biologically significant or with greater univariate significance to the dependent variable. Multicollinearity was checked by calculating the variance inflation factor (VIF) function (R package *car* version 3.1-3) (Fox and Weisberg 2019) and comparing correlation plots (R package *PerformanceAnalytics* version 2.0.8) (Peterson and Carl 2024). For example, high correlation was observed between distance to PCR deer positives, distance to Michigan, and deer habitat. Distance to deer PCR positives, being the primary variable of interest, was retained in the model over distance to Michigan and deer habitat. The final full model included five variables: (1) Distance to 17 PCR positive deer (2021); (2) Deer harvest in the fall; (3) average COVID-19 cases reported in the fall by public health unit by year; (4) average human population by year; and (5) year of sample (Table 1). Some variables were first rescaled by 10 or 1000 to allow the model to converge.

For the GLMM models, samples with missing data for key variables were removed (i.e. samples in 2024 where data was not yet available for COVID-19 cases and fall deer harvest). The one deer that tested PCR positive but seronegative was retained in the serology data as a “negative”, as this wouldn’t bias the results towards rejecting the null hypothesis that there was no association between seropositivity and the landscape. The model was split into training and testing data (20/80%; N=195/785). Rather than using an automated stepwise approach, which is not recommended in GLMMs, model selection was driven by comparing the AIC of multiple models, a measure of goodness of fit that penalizes overfitting. Using the *drop1* function of package *lme4* (Bates et al. 2015), five new models were created from the full model using the training data, each with just one variable was dropped. The model with the lowest AIC value was retained and compared to the previous model using an ANOVA test. If the model reduced the AIC and the ANOVA test was non-significant, this model was retained and the *drop1* process repeated until the most parsimonious model remained. All figures were generated with R package *ggplot2* (Wickham 2016), and all code and non-restricted datasets will be made available on OSF upon publication.

## Results

### SARS-CoV-2 surveillance in southern Ontario

Between January 2022 and June 2024, 2,839 animals were tested for current or past infection of SARS-CoV-2 across southern Ontario using serological or molecular analyses. Only animals with location data were retained in the sample. The study area covered 148,460 km², with SARS-CoV-2 infection detected in wildlife across 40,960 km² (28% of the study area). Among the 15 projects that sourced samples between 2022 and 2024 (Table 2), 39.0% (N=1107) were deer harvested through the CWD program run by MNR, followed by animals intentionally live-trapped for SARS-CoV-2 or sampled through the MNR Rabies Program TVR project (N=482; 17%), animals sampled at wildlife rehabilitators (N=327; 11.5%), and *Peromyscus* sampled at Algonquin Provincial Park (N =211; 7.4%). Overall, 76% of the animals sampled at wildlife rehabilitators were orphaned juveniles. Surveillance of roadkill accounted for 5.2% of samples collected (N=148). Between 2021 and 2023, 35 mink and 57 other furbearer carcasses (beaver, fisher, river otter) were provided by local trappers for testing (N=92; 3.2%). The remaining samples were collected on assorted ecology projects run by MNR or Trent University, or ungulates submitted through the Canadian Wildlife Health Cooperative (CWHC).

**Table 2.** Breakdown of samples by project 2022-2024.

| Project | 2022 | 2023 | 2024 | Total | % All samples |
| --- | --- | --- | --- | --- | --- |
| Chronic Wasting Disease surveillance | 626 | 481 |  | 1107 | 39.0% |
| Wildlife in rehabilitation | 156 | 160 | 11 | 327 | 11.5% |
| SARS-CoV-2 surveillance live-trapping (2023) |  | 288 |  | 288 | 10.1% |
| Algonquin Park Small Mammal Project | 75 | 136 |  | 211 | 7.4% |
| SARS-CoV-2 surveillance live-trapping (2022) | 156 |  |  | 156 | 5.5% |
| Roadkill, hunted or euthanized (not CWD) | 47 | 85 | 16 | 148 | 5.2% |
| White-tailed deer live-trapping | 96 | 44 |  | 140 | 4.9% |
| Rabies dRIT laboratory testing | 133 |  |  | 133 | 4.7% |
| Zoo animals | 87 | 23 | 8 | 118 | 4.2% |
| Mustelid and furbearer carcass samples | 86 | 6 |  | 92 | 3.2% |
| Ungulates tested by CWHC | 23 | 27 | 7 | 57 | 2.0% |
| Rabies TVR project | 5 | 23 |  | 28 | 1.0% |
| Eastern coyote and Algonquin wolf live-trapping | 13 |  | 1 | 14 | 0.5% |
| Highly Pathogenic Avian Influenza outbreak response live-trapping (February 2024) |  |  | 10 | 10 | 0.4% |
| Wild turkey live-trapping | 10 |  |  | 10 | 0.4% |
| **Total** | **1513** | **1273** | **53** | **2839** | **100%** |

Overall, 95.7% of the samples since 2022 were native wildlife species (Table 3 and 4). White-tailed deer were the most common species sampled (N = 1446; 50.9%). Among all cervids, the majority were sourced through the CWD Program (1107; 72.2%), followed by live-trapping (N=140; 9.1%), roadkill or hunted (N=84; 5.5%), zoos (N=76; 5.0%), wildlife rehabilitators (N=69; 4.5%), and carcasses submitted to CWHC (N=57; 3.7%). Other frequently sampled species were deer mice (N=211; 7.4%), raccoons (N=192; 6.8%), and striped skunks (N=155; 5.5). Sixty-eight mink were sampled (2.4%) between 2022 and 2024. Most animals were male (50.1%, 40.5% female, 9.4% unknown) and adult (69.5%) (26.7% juveniles, 3.8% unknown).

**Table 3.** Summary of the serological tests by species and species class. In 2021, two deer were both seropositive and PCR-positive.

| Total Samples (# Seropositive) / % |  |  |  |  |  |  |  | Combined<br>Sample<br>2019-2024) | Combined<br>Sample<br>(2022-2024) |
| --- | --- | --- | --- | --- | --- | --- | --- | --- | --- |
|  | Species | 2019 | 2020 | 2021 | 2022 | 2023 | 2024 |  |  |
| NATIVE | White-tailed deer<br>(*Odocoileus virginianus*) | 50 (0) | 160 (1)<br>1% | 89 (6)<br>7% | 463 (51)<br>11% | 517 (36)<br>7% | 11 (2)<br>18% | 1290 (96)<br>7% | 991 (89)<br>9% |
|  | Raccoon (*P. lotor*) |  | 140 (0) | 135 (0) | 10 (0) | 77 (0) | 3 (0) | 365 (0) | 90 (0) |
|  | Virginia opossum (*D. virginiana*) |  |  |  | 33 (1)<br>3% | 65 (0) | 7 (1)<br>14% | 105 (2)<br>2% | 105 (2)<br>2% |
|  | American mink (*N. vison*) |  |  | 52 (0) | 37 (0) | 12 (0) | 3 (1)<br>33% | 104 (1)<br>1% | 52 (1)<br>2% |
|  | Striped skunk (*M. mephitis*) |  | 36 (0) | 19 (0) | 18 (0) | 14 (0) |  | 87 (0) | 32 (0) |
|  | North American beaver (*C. canadensis*) |  |  |  | 39 (0) |  |  | 39 (0) | 39 (0) |
|  | Eastern coyote (*Canis latrans*) or<br>Algonquin wolf (*C. lupus lycaon*) |  |  |  | 13 (0) | 1 (0) | 1 (0) | 15 (0) | 15 (0) |
|  | North American river otter<br>(*L. canadensis*) |  |  |  | 11 (1)<br>9% |  |  | 11 (1)<br>9% | 11 (1)<br>9% |
|  | Wild turkey (*M. gallopavo*) |  |  |  | 10 (0) |  |  | 10 (0) | 10 (0) |
|  | Fisher (*P. pennanti*) |  |  |  | 6 (0) | 1 (0) |  | 7 (0) | 7 (0) |
|  | Red fox (*V. vulpes*) |  |  |  | 1 (0) | 2 (0) |  | 3 (0) | 3 (0) |
|  | White-footed mouse (*P. leucopus*) |  |  |  |  | 2 (0) |  | 2 (0) | 2 (0) |
|  | Long-tailed weasel (*Mustela frenata*) |  |  |  |  | 1 (0) |  | 1 (0) | 1 (0) |
|  | American red squirrel<br>(*Tamiasciurus hudsonicus*) |  |  |  |  | 1 (0) |  | 1 (0) | 1 (0) |
| EXOTIC | Barbary sheep (*Ammotragus lervia*) |  |  |  |  |  | 5 (0) | 5 (0) | 5 (0) |
|  | Eurasian wild boar (*S. scrofa*) |  |  |  |  |  | 2 (0) | 2 (0) | 2 (0) |
|  | Porcupine caribou<br>(*Rangifer tarandus granti*) |  |  |  |  |  | 2 (0) | 2 (0) | 2 (0) |
|  | Linnaeus's two-toed sloth (*Choloepus didactylus*) |  |  |  |  |  | 1 (0) | 1 (0) | 1 (0) |
| Total |  | 50 (0) | 336 (1)<br>0% | 295 (6)<br>2% | 641 (53)<br>8% | 693 (36)<br>5% | 35 (4)<br>11% | 2050 (100)<br>5% | 1369 (93)<br>7% |

**Table 4.**
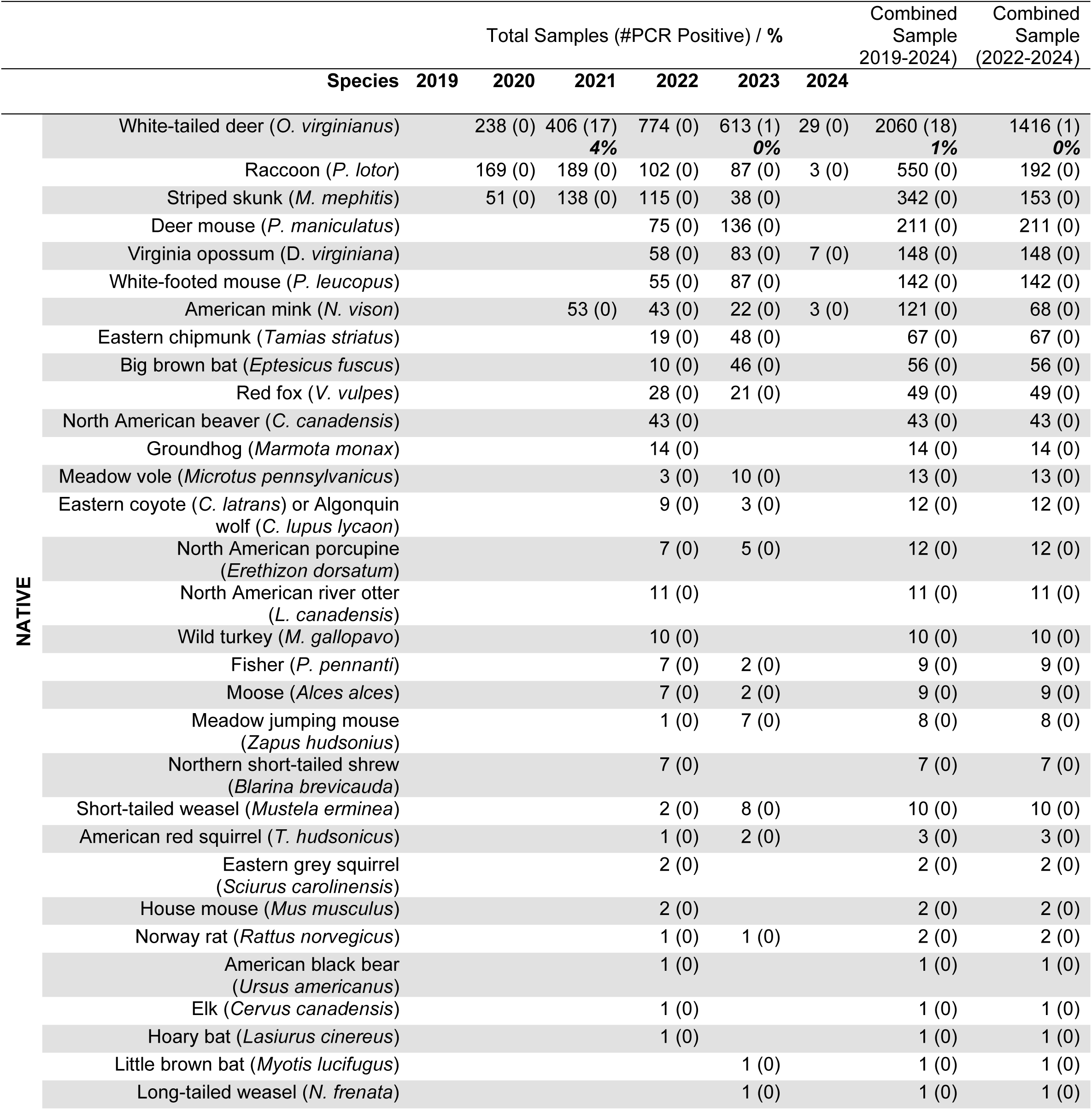
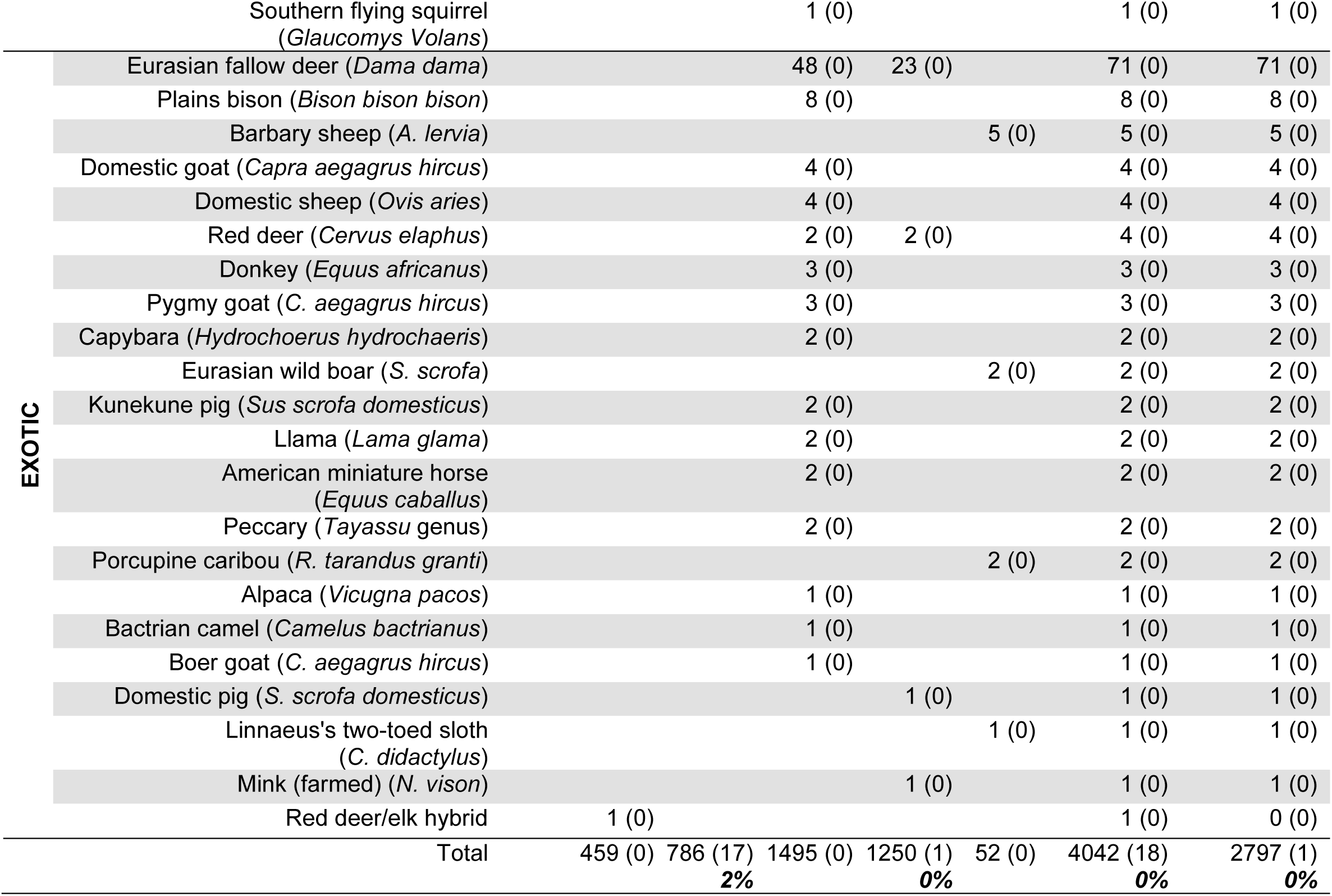
Summary of PCR tests by species and species class. In 2021, two deer were both seropositive and PCR-positive. The 17 positive deer in 2021 were those previously reported by Pickering et al. (2022).

### Serology and PCR results

Between January 2022 and June 2024, 93 animals tested positive for SARS-CoV-2 antibodies (Table 3), and viral RNA was detected through PCR testing in one deer in central Ontario in November 2023 (Table 4). The PCR-positive deer was seronegative and located over 235 km away from nearest PCR-positive deer case from 2021 (Figure 1). The majority of seropositive animals were detected in 2022 (N= 53; 57.0%), followed by 2023 (N= 36; 38.7%) and 2024 (N=4 ; 4.3%). Among the 93 seropositive animals, 89 were deer, and the remaining included new species detections for the region: one river otter (March 2022), two Virginia opossum (October 2022, February 2024), and an American mink (February 2024) (Figure 2). Only one animal sampled during live-trapping tested seropositive, a Virginia opossum sampled in February 2024 near Windsor, Ontario, close to the Michigan/Ontario border. The two serologically positive cases in February 2024 (mink, opossum) were separated by 480 km.

**Figure 1.**
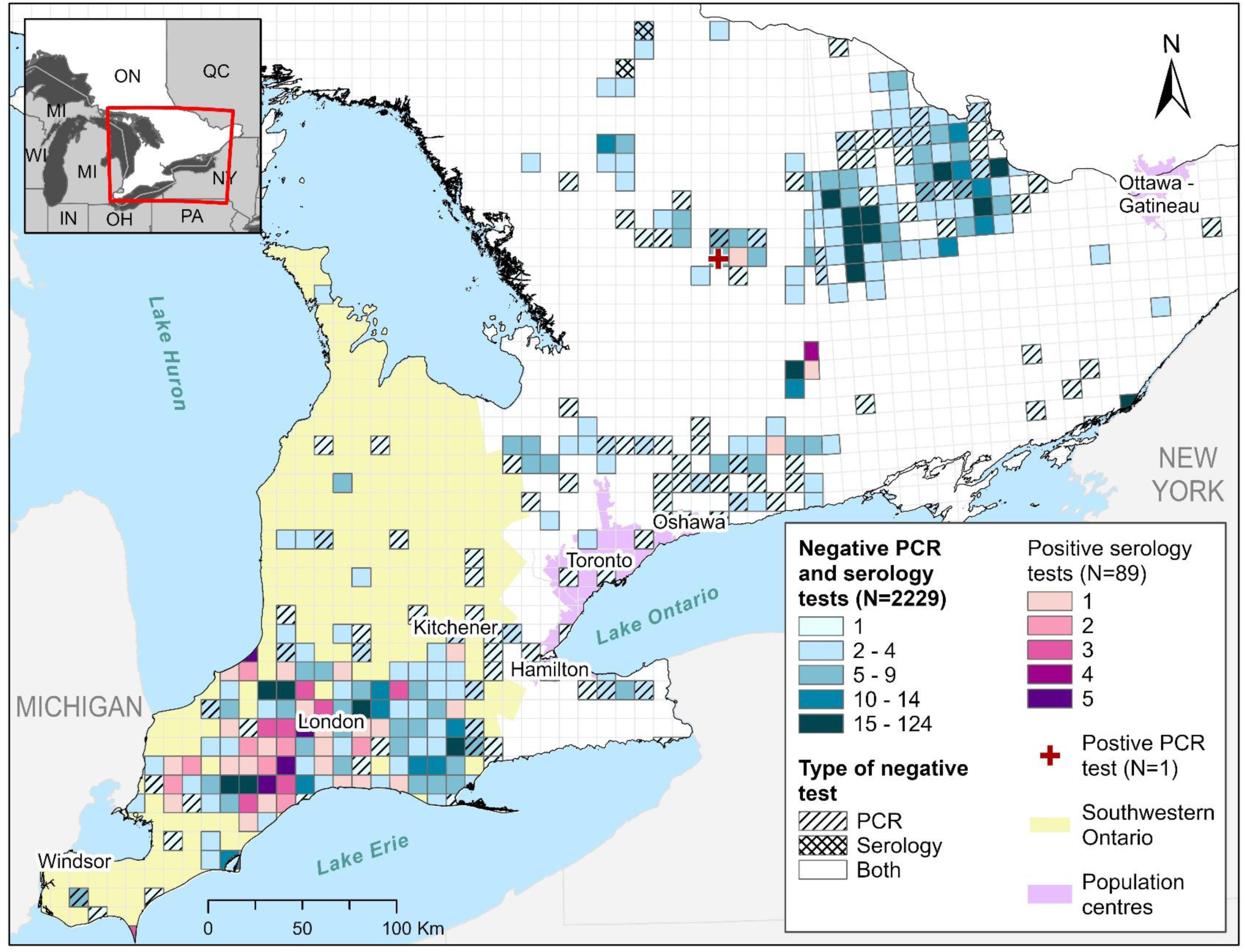
Distribution of SARS-CoV-2 positive deer (89 seropositive and one PCR-positive deer; ID= wcov1242165) relative to the sampling effort among deer only within each 10 x 10 km grid cell. Provincial and state borders were sourced by Statistics Canada (Statistics Canada 2023c) and the U.S. Census Bureau (U.S. Census Bureau 2018).

**Figure 2.**
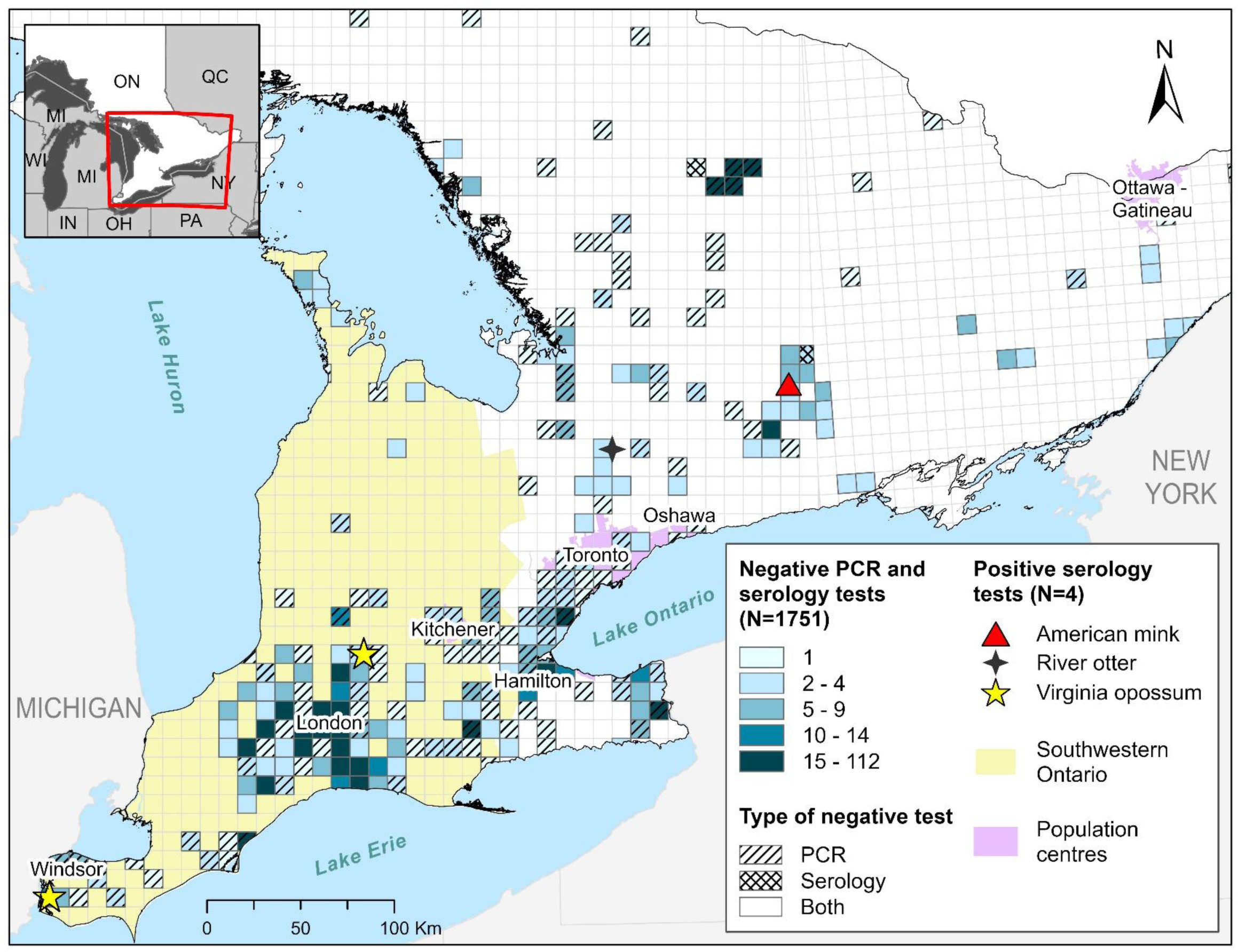
Distribution of four SARS-CoV-2 serological positives mammals, excluding deer, relative to the sampling effort of these species within each 10 x 10 km grid cell. Provincial and state borders were sourced by Statistics Canada (Statistics Canada 2023c) and the U.S. Census Bureau (U.S. Census Bureau 2018).

The PCR-positive deer was detected during annual CWD surveillance. The CWD project accounted for 79.6% (N=74) of the serologically positive animal cases, followed by roadkill or juvenile deer collected during CWD sampling (12.9%; N=12), live-deer trapping and collaring (N=5; 5.3%), collected through furbearer trapping (1.1%; N=1), and during Avian influenza sampling (1.1%; N=1). No animals from rabies surveillance, *Peromyscus* live*-*trapping, canid live-trapping, ungulates submitted by CWHC, or any captive facilities were PCR positive or seropositive.

### Sequence analysis

A high-quality genome sequence was assembled for the one PCR-positive deer (sample wcov1242165) following sequencing and bioinformatics analysis with 99.6% genome coverage at 10X or greater depth of coverage and 23,235X mean coverage depth. Both UShER and Nextclade assigned wcov1242165 to the GS.3 PANGO lineage, alias of recombinant lineage XBB.2.3.11.3, first detected in the USA in May 2023 according to GISAID (Elbe and Buckland-Merrett 2017). Compared to Wuhan-Hu-1, the wcov1242165 genome had 33 synonymous nucleotide substitutions and 75 amino acid substitutions, as well as 4 inframe deletions affecting ORF1a, S and N gene amino acid sequences (ORF1a:del3675/3677, S:del25/27, S:del144, N:del31/33) commonly found in lineage GS.3 sequences according to outbreak.info (Gangavarapu et al., 2023). The wcov1242165 genome had several amino acid mutations not typically found in GS.3 sequences (ORF1a:T403I, ORF1a:P3371S, ORF1a:L3588F, ORF1a:A4092V, ORF1b:E1601D, S:A27S, S:G72E, ORF9b:P10S) according to outbreak.info. Of these mutations, ORF1ab:A4092V, S:G72E and 6 nucleotide mutations (T3232C, C12540T, C19113T, G21777A, C29659T, C29719T) were unique to wcov1242165 compared to nearest neighbour sequences from UShER phylogenetic placement analysis against 16,884,360 genomes from GISAID, GenBank, COG-UK and CNCB (as of 2025-03-26) and the sarscov2phylo 13-11-20 tree with newer sequences added by UShER.

Phylogenetic analysis recovered the new wcov1242165 sequence from deer deeply nested within a clade of SARS-CoV-2 sequences from humans (Figure 3, A-B). The most closely related viral sequences were from humans sampled in Ohio and Texas, USA, and Ontario, Canada, in 2023. Phylogenetic analysis revealed that the new wcov1242165 genome from deer was distinct from other publicly available deer-derived SARS-CoV-2 sequences, which were not recovered in the same cluster of human-derived SARS-CoV-2 as the new sequence from deer.

**Figure 3.**
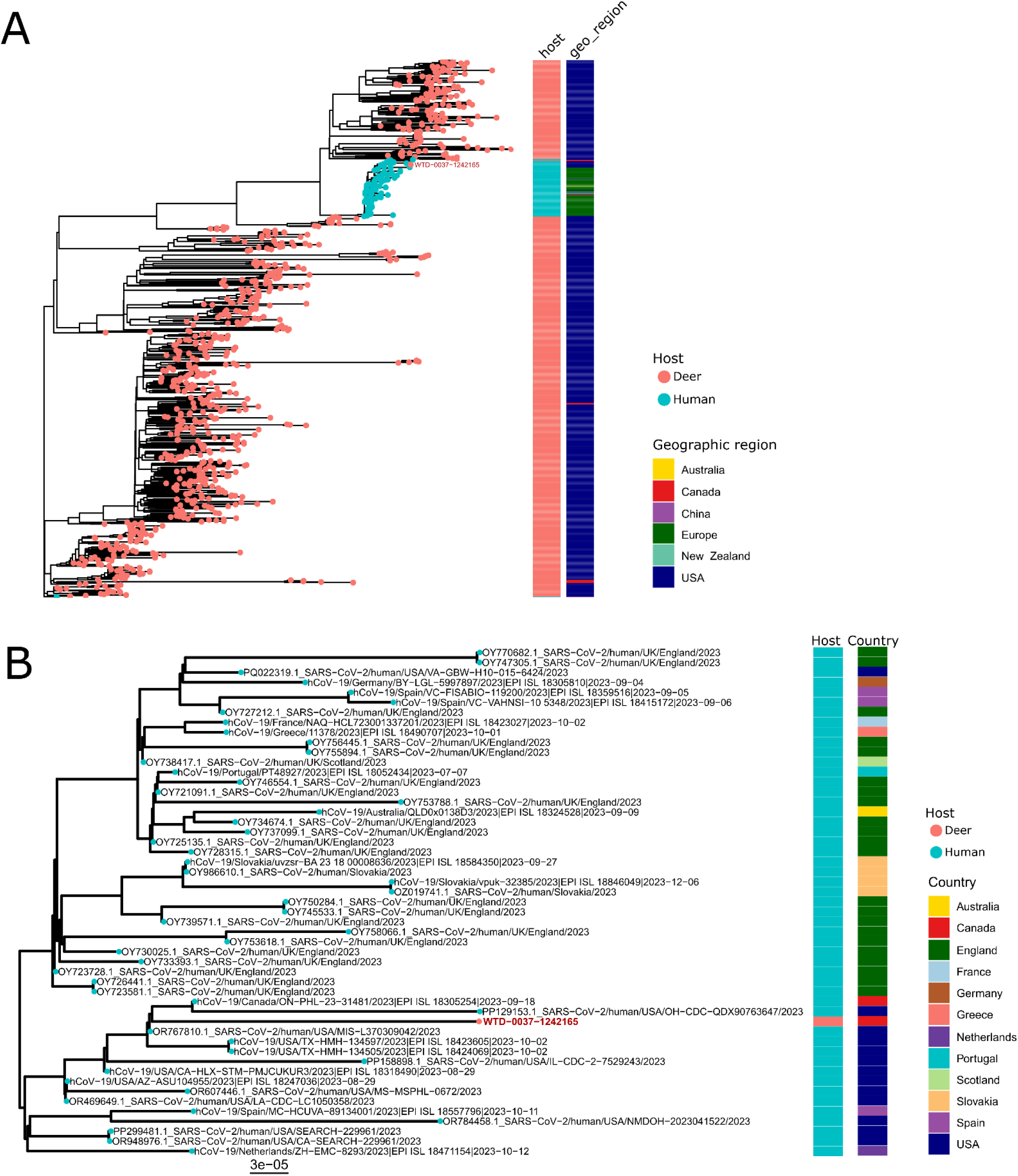
Maximum likelihood phylogeny of SARS-CoV-2 genomes from white-tailed deer. (A) Phylogeny showing all publicly available SARS-CoV-2 genomes from deer (N=811) and (B) 96 closest matches to the WTD-0037-1242165 identified by UShER and a close-up of a subtree showing placement of the new viral sequence from deer deeply nested within a group of viral sequences from human hosts. Tips of the phylogeny are coloured according to the host type (human or deer). The new viral sequence from deer is labeled in red font.

### Temporal and spatial patterns in SARS-CoV-2 detections

The majority (90.3%) of seropositive animals since 2022 were found in the region of Southwestern Ontario (Figure 1 and 2). The seroprevalence among deer tested for antibodies was high (15.2%) in this region. Among Virginia opossums, the only other positive animal in the southwest, the prevalence was 2.0%. Overall, 10.5% (N=84) of animals tested for antibodies in Southwestern Ontario were seropositive.

Prior to 2022, 7 deer (2.3% of deer sampled for antibodies) across the whole study area were seropositive, compared to 89 deer during or after 2022 (9.0%) (Tables 3). Between 2019 and 2021, only deer and raccoons were tested, so we don’t have a time comparison for all species. Overall, prior to 2022, 1.0% of all animals tested for antibodies were seropositive, compared to 6.8% since 2022.

We compared the prevalence of SARS-CoV-2 PCR and seropositive cases from 2019-2024 against reported human COVID-19 cases reported each fall (September - December) in the same public health units where the samples were collected (Ontario Agency for Health Protection and Promotion (Public Health Ontario) 2024). No human COVID-19 data were available for fall of 2024, as sample collection ended in June 2024. The proportion of seropositive tests in this study increased from 0.3% in 2020 to 11.4% 2024, with a drop in 2023 (5.2%). Although seroprevalence was highest in 2024, only 35 serological tests were carried out this year. Just by observation, the peak activity in human cases occurred in the fall in 2021, coinciding with the highest yearly proportion of the samples that tested PCR positive. Differences in seropositivity by sex and age were compared across the whole serological dataset (2022-2024; N =2,839) among deer tested for antibodies using Fisher’s exact test. Sex and age were not significantly associated with seropositivity (p>0.05).

### Regression modeling

#### Model selection

The most parsimonious model (lowest AIC through stepwise selection) explaining seropositivity of deer included three variables retained in the model, which were all significant with presence of SARS-CoV-2 on the landscape (distance to PCR positive deer, fall deer harvest, and human population) (Table 5). Distance to the previous 17 PCR positive deer was the most significant variable (p=0.0008), with every 10 km distance away from a previous positive PCR case, or the equivalent of one grid 10 x 10 km cell, the odds of an animal being seropositive decreased by 7.6% (Odds Ratio (OR): 0.92; 95% CI: 0.88 – 0.97).

**Table 5.** GLMM model coefficients of final model.

|  | Estimate | Std. Error | Z value | P Value |
| --- | --- | --- | --- | --- |
| **Intercept** | -3.6508 | 0.7240 | -5.0422 | <0.001 |
| **Distance to 17 PCR+ deer (2021)** | -0.0790 | 0.0235 | -3.3598 | <0.001 |
| **Annual fall deer harvest (per 1000 animals)** | 0.4890 | 0.2034 | 2.4045 | 0.0162 |
| **Human population (per 1000 people)** | 1.1641 | 0.4172 | 2.7906 | 0.0053 |

The annual fall deer harvest was positively correlated with seropositivity, where deer sampled in grid cells with higher average hunting numbers were more likely to be seropositive. For every 1000 deer harvested in a WMU, the odds of one animal being seropositive increased by 63% (OR: 1.63; 95% CI: 1.1 – 2.5). Confidence intervals were calculated using the R *Stats* package (R Core Team, 2023). Population was found to be significant in the multivariate model, but not in univariate analysis. Population was not highly correlated with other variables, so this result was not due to multicollinearity. The results indicate that for every 1000 people, the odds a deer tested seropositive increased three times (OR: 3.2; 95% CI: 1.4 – 7.8).

#### Model diagnostics

Diagnostics were carried out of the mixed model using R package *PerformanceAnalytics* and *DHARMa*, which was created for assessing GLMMs by simulating standardized residuals (Hartig 2024). Dharma plots of residual vs predicted values showed no significant problems and no evidence of significant outliers, overdispersion or underdispersion. In GLMMs, the models assume a linear relationship between continuous predictor variables and the logit of the outcome. We ran scatterplots and found a linear trend with deer positives and human population, but some violation for deer harvest, which is not surprising with ecological models.

#### Model performance and accuracy

Model performance of the final model was assessed by making predictions with the test data and determining the model’s accuracy with R package *lme4* (Bates et al. 2015). The *Predict* function uses population level fixed effects, where the corresponding random effect components are set to zero.

To test the ability of this model to correctly classify samples as positive or negative, we identified an optimal threshold of 0.1, which maximizes sensitivity and specificity. This threshold was used to create an ROC curve using R package *pROC* (Robin et al. 2011), where the area under the curve (AUC) represents the probability that a model will correctly distinguish between true positives and true Negatives, where a perfect score of 1 indicates perfect separability. The model had an AUC value of 0.71, which is considered acceptable discrimination (Mandrekar 2010). The model has low sensitivity (60%), but moderate specificity (88%), indicating lower ability to detect true positives (i.e., a higher false negative rate).

## Discussion

We initiated an active surveillance program following the discovery in 2021 of a divergent lineage of SARS-CoV-2 in deer. Analyses of B.1.641 suggested that the lineage may have evolved in deer for up to one year, accumulating 76 mutations compared to ancestral SARS-CoV-2 and appearing to have originated in Michigan mink farms (Pickering et al. 2022). Despite sampling 1,446 Ontario deer during our active surveillance program since 2022, we have not detected additional B.1.641 infections, or infections with related B.1 viruses. The only SARS-CoV-2 infection we have observed in deer from Ontario after 2021 is a single XBB infection in central Ontario. Consequently, our data do not support the conclusion that deer in Ontario are a reservoir of B.1.641 in particular, or SARS-CoV-2 in general.

We define a reservoir as having an ability to maintain infection with a zoonotic pathogen, and the presence of an interface for transmission to humans (Haydon et al. 2002). While it is clear that there is ample opportunity for transmission of SARS-CoV-2 from deer to humans, such as through hunting and backyard feeding of deer (Pratt et al. 2025), the lack of any viruses related to B.1.641 in our study suggests that the virus has not been maintained in the deer population following the relatively localised year-long outbreak previously observed (Pickering et al. 2022). Instead, subsequent infections (e.g., the XBB infection we observed) are likely a result of additional spillovers from humans. A similar pattern of multiple spillover events has been observed (Kuchipudi et al. 2022; Caserta et al. 2023), including detection of Omicron variants at a time where it accounted for 90% of human cases (Vandegrift et al. 2022), although there is also evidence of sustained transmission within deer (Caserta et al. 2023; Feng et al. 2023). However, we consider that evidence is accumulating which indicates that deer populations do not sustain SARS-CoV-2 infections indefinitely.

Across Ontario, we estimated that SARS-CoV-2 seropositivity among deer was about 9.0%, but in Southwestern Ontario, the estimated seropositivity was 15.2% (Figure 1 and 2). It is clear that there was some transmission of B.1.641 among deer, both because the lineage appeared to evolve for about a year prior to its detection in 2021 and because the best predictor in our regression model for a seropositive deer was the distance to a previous B.1.641 case. This variable in combination with variables describing human population density and deer harvest density allowed us to estimate that for every 10 km distance away from a previous B.1.641 case, the odds of a deer being seropositive decreased by 7.6%. The regression model paints a picture of B.1.641 spreading among deer for some period of time in areas with high deer and human population densities. The infections eventually ceased to spread however, and this lack of sustained transmission may be due to patterns in contact between animals or the adaptive immune response of deer. The decrease in SARS-CoV-2 in deer and the parallel increase in seropositivity raises the question of whether previous infection provides protective immunity, which may explain why prevalence of infection decreased from 2021 to 2024. Although titres will drop with time, neutralizing antibodies have been found to persist in captive white-tailed deer for at least 13 months (Hamer et al. 2022).There is much that we do not yet know about how the deer immune system responds to SARS-CoV-2 infections (Kotwa et al. 2023) or how anthropogenic and natural landscape barriers that influence deer dispersal consequently affect disease transmission (Peterson et al. 2017).

In addition to the seropositive deer detected in proximity to known B.1.641 infections, we also detected a small number of other seropositive species (two opossums, a mink, and an otter). With the exception of one opossum, none of the positive species occurred in close proximity to known B.1.641 cases, and as such, we have no evidence to suggest B.1.641 was spilling from deer into other wildlife species. In general, the seroprevalence was low in non-deer species, which may reflect independent spillover events to these wildlife species without onward and sustained intra-species transmission. Low estimates of seropositivity might also be due to a lack of seroconversion, or waning antibodies. For example, an opossum resampled after a month had a low percent of neutralizing antibodies (51.1%; Goldberg et al., 2024). While the Surrogate Virus Neutralization Test (sVNT) used to detect neutralization antibodies has been highly sensitive (95-100%) to Covid-19 in humans (Tan et al. 2020), this may vary based on species, method of testing and field conditions, and even mutations since this test was first established (Hewitt et al. 2025). For example, there is some evidence that Nobuto strips may underestimate seroprevalence for pre-Omicron variants (Hewitt et al. 2025). Another factor that cannot be accounted for is the potential cross-reactions with other endemic coronaviruses that could be affecting the specificity of the assay. Little is known about endemic coronaviruses circulating in these species. As for low prevalence of virus detected through PCR testing, it is possible that active infection was missed due to the means of swabbing. In experimentally inoculated skunks, viral titres were greater in nasal samples compared to oral samples (Bosco-Lauth et al. 2021), the latter of which were primarily used in our study, although this exposure to high titred innoculum does not reflect natural infection pressure.

Human SARS-CoV-2 cases peaked in Ontario during 2021, and not surprisingly this was also the peak of SARS-CoV-2 infections in Ontario’s wildlife. Superficially, this appears to support the idea that high infection burden in the human population was associated with spillover to wildlife (e.g., Vandegrift et al., 2022). All of the documented positive cases of SARS-CoV-2 in wild animals during 2021 were deer with B.1.641. However, as of April 2021, Delta became the dominating circulating variant in the human population (Ontario Agency for Health Protection and Promotion (Public Health Ontario) 2022), and we have not documented any Delta infections among wildlife in Ontario. The B.1.641 infections appear to have been evolving in deer for up to a year (Pickering et al. 2022). We did observe a peak in seropositivity among wildlife samples in 2022, a year after the peak in B.1.641 cases, mostly, but not exclusively arising from deer samples. We cannot disentangle whether the seropositive cases represent spillovers from the human population, or sylvatic transmission among wildlife species.

As has been observed in other North American jurisdictions since Omicron became the dominant SARS-CoV-2 clade (Feng et al. 2023; McBride et al. 2023), we found recent evidence of a deer with an Omicron infection (Figure 3). This does not appear associated with the previous B.1.641 infections, but is instead likely a recent spillover from the human population. The infected deer was sampled in central Ontario in a region distant from any other documented cases of deer with SARS-CoV-2 and with an abundance of cottages, providing opportunities for human-to-deer transmission. The lack of additional cases in deer is likely associated with the reduced prevalence of infection in the human population after 2021. It is also possible, however, that deer are less susceptible to infection with the Omicron lineages of SARS-CoV-2 than with older lineages. One possible explanation for such an effect is that APOBEC family of proteins, which lead to C>U mutations through RNA editing, are prominent features of deer immune response to infections with pre-Omicron lineages (Pickering et al. 2022; Kotwa et al. 2023). APOBECs have also been shown however, to promote viral replication and propagation (Kim et al. 2022). Therefore, if APOBEC-mediated RNA editing is less involved in deer immune response to Omicron, this might reduce prevalence of infection among deer. There remains considerable uncertainty about the differential immune response of deer to lineages of SARS-CoV-2.

Filling the knowledge gaps in viral genomic surveillance and the interface between human and animal disease can help us better prepare for emerging diseases and can improve our understanding of host jumping (Mubareka et al. 2023; C.C.S. Tan et al. 2024). When disease surveillance systems exist across animal and human health sectors however, they are often carried out by separate agencies or industries with different regulations and reporting requirements, making communication and data sharing difficult. To address these barriers, we applied a One Health approach to our surveillance program, with the aim to reduce the existing gaps separating huma and animal disease knowledge. We established a multidisciplinary team, with cross-sectoral expertise in animal, human, and environmental health (Lee et al. 2024). This follows Kuchipudi et al.’s call for a robust One Health-centered SARS-CoV-2 surveillance of cross-species transmission in non-human species (Kuchipudi et al. 2023).

We recognize that our study has potential limitations. For example, on live wild animals, we principally used oral swabs for virus detection, and this may not have been the optimal virus target in some species. Our serological assay was not validated for use across all species in the study. We also need to establish a better understanding of the immune response of deer which might have led to the eventual decline of B.1.641, and perhaps to less prevalent Omicron infections. With limited data on serological and viral infection, sex and age-mediated differences could be missed, especially when 7% of the animals were of unknown sex, which may have limited the influence on transmission and behavior.

Wildlife reservoirs of SARS-CoV-2 can alter virus ecology, leading to divergent lineages with potential health consequences for the human population. Effects of the B.1.641 lineage on humans are not yet well understood, and consequently there has been considerable interest in assessing the possibility that deer were becoming a reservoir of this virus. We established a multi-disciplinary team to undertake a One Health-centred surveillance program to evaluate the risks of reservoir establishment. To date, evidence suggests that B.1641 has not continued to circulate among deer, and therefore, we have no evidence that deer are a reservoir of SARS-CoV-2 in Ontario.

## Acknowledgements

We want to thank those who made sampling wildlife possible, including students and volunteers assisting on the SARS-CoV-2 project (Laurelie, Brennan, Alexis, Sean, Thomas, Greg, Sarah, Maissie, Shelby, Sarah, Rebekah, Janet), MNR researchers who took on additional sampling work to increase our study area, as well as wildlife rehabilitators (Another Chance, Aspen Valley Wildlife Sanctuary, Erie Wildlife Rescue, Fur-Ever Wild Rehabilitation, Hobbitstee Wildlife Refuge, Itty Bitty Critters Wildlife Rehabilitation, Procyon Wildlife, Salthaven Wildlife Rehabilitation & Education Centre, Shades of Hope Wildlife Refuge, Woodlands Wildlife Sanctuary, Wings Rehabilitation Centre, with special thanks to Carol, Colleen, Kelly, and Dave), zoos (Greenview Park & Zoo, Marineland, Waterford Deer Park, West Perth Animal Park, Riverview Park & Zoo), Conservation Authorities (Ausable Bayfield CA, Catfish Creek CA, Kettle Creek CA, Lower Thames Valley CA, Upper Thames River CA, St Clair Region CA), provincial and national parks (Algonquin, Komoka, Pinery, Point Pelee, Short Hills, Rondeau), the CWHC (B. Stevens and L. Shirose), and municipalities (City of London, City of St. Thomas, Town of St. Mary’s, and Oxford County). Those who contributed harvested deer samples, we thank members of the Haudenosaunee Confederacy and Haudenosaunee Wildlife and Habitat Authority, Kettle and Stony Point First Nation, licenced Ontario deer hunters, and Appin BBQ and Catering. We also thank Norfolk Field Naturalists and the Niagara SPCA andHumane Society.

We recognize that our study area in present-day Ontario is located on lands of traditional multiple Indigenous nations, and that accessing wildlife on these lands has been possible in part by the preservation of greenspaces across Ontario and through stewardships with these Indigenous communities.

## Author contributions

Conceptualization: L.C., J.D.K., O.L., F.M., B.,P., C.J., S.M., J.B.

Data Curation: L.C., J.D.K., S.P.J., C.L., E.C., W.Y., P.K., O.V., O.L., A.I.S., C.J., H.C., S.M., J.B.

Formal Analysis: L.C., J.D.K., S.P.J., C.L., A.D., P.K., O.V., H.C., S.M., J.B.

Funding Acquisition: O.L., A.I.S., B.,P., C.J., S.M., J.B.

Investigation: L.C., J.D.K., S.P.J., C.L., A.D., A.S., S.L.N., E.C., W.Y., H.C., J.B.

Methodology: L.C., J.D.K., C.L., A.D., P.K., O.V., O.L., S.M., J.B.

Project Administration: J.D.K., C.J., H.C., S.M., J.B.

Resources: J.D.K., O.L., C.J., H.C., S.M., J.B. Software: L.C., J.D.K., P.K., O.V.

Supervision: L.C., J.D.K., O.L., C.J., H.C., S.M., J.B.

Validation: J.D.K., C.L., A.D.

Visualization: L.C., P.K., O.V.

Writing – Original Draft Preparation: L.C., J.D.K., S.P.J., W.Y., O.V., O.L., H.C., J.B.,

Writing – Review & Editing: L.C., J.D.K., S.P.J., C.L., A.D., A.S., S.L.N., E.C., W.Y., P.K., O.V., O.L., A.I.S., F.M., B.,P., C.J., H.C., S.M., J.B.

## Declaration of interests

The authors declare no competing interests.

## Notes

### Competing Interest Statement

The authors have declared no competing interest.

### Summary of Updates

An author was added to the manuscript; Table 3 and 4 revised.

